# Threading the needle: Spatial constraints sharpen visual sensitivity in honeybees

**DOI:** 10.64898/2026.01.02.696520

**Authors:** T. Jakobi, M. Garratt, M. Srinivasanand, S. Ravi

## Abstract

Navigating dense clutter requires honeybees to execute rapid visuomotor comparisons to identify the best route. How they resolve perceptual inputs to make a passage choice from multiple options is an open question. We studied this in a free flying paradigm where bees were challenged to choose between a taller and shorter aperture with varying height difference. We monitored choice across four height difference ratios at three spatial scales. Bees chose the taller aperture in all tests, but the precision of these choices depended heavily on the absolute size of the gap. For each size scale of aperture pairs, psychometric modelling revealed that when making a choice bees generally evaluate the relative height difference, conforming to Weber’s law. However, bees’ choices were non-uniformly sensitive across aperture size scales. Bees had greater sensitivity at the smallest spatial scale, where the likelihood of collision danger is larger. This deviation in behaviour due to absolute aperture size suggests a dynamic risk-cost trade-off. Bees appear to prioritise costly, high-acuity inspections only when imposed by critical physical constraints of the environment, relaxing this vigilance at safer, larger scales to conserve energy.

## Introduction

Agile navigation through cluttered spaces is critical to the survival and foraging success of flying insects. Honeybees, with their short interocular separation, rely on (what is effectively) monocular vision to select viable routes from a large number of options, often with marginal geometric advantages [1–4], despite resolution constraints. These decisions demand rapid visual assessment and efficient processing to minimise collision risks and optimise paths amid ecological pressures and resource limitations [2,5].

In many organisms (including humans), the sensory percept of a change in a stimulus is not proportional to the magnitude of the absolute change; rather, it is proportional to the *relative* change in the stimulus. This principle is known as Weber’s law, which states that the difference threshold is a constant fraction of the stimulus magnitude [6]. This leads to a logarithmic scaling in sensation strength, as elaborated in extensions like Fechner’s law [7]. This phenomenon has been observed and documented in relation to several human modalities—for example, the perception of light, sound and weight. This phenomenon has also been observed in honeybees, in relation to the perception of object size [8], colour [9], numerosity [10], and even sugar solution concentration [11].

Honeybees also discriminate the sizes of passages during spontaneous navigation through clutter to determine the safer path, selecting wider gaps in dual-aperture scenarios down to a pair width difference of 2 cm (67% wider) [12]. This preference scales with relative width, following Weber’s law, and causing a typical sigmoidal logistic curve as perception saturates at large proportional differences [6]. This visual sizing is mediated by the optic flow (OF) generated by the gap edges during the insect’s approach [13,14]. In principle, this motion-based cue for size could provide far finer discrimination acuity than what has been tested in existing choice-based paradigms. Evidence for such precision is documented in bumblebees, which modulate their body orientation by minute fractions to minimise their profile when traversing narrow gaps [15]. This suggests that the full perceptual limits and cognitive strategies governing route selection and rapid decision-making are underexplored, warranting further investigation.

Roaming animals encounter speed-accuracy trade-offs (SATs) in their environments, balancing rapid responses against precision to optimise behaviour under time and risk constraints [16]. In bumblebees, SATs are manifested in foraging, where speed is sacrificed for accuracy because errors are penalised (for example, when choosing unrewarding flowers) [17]. During flight-based tasks, such as inspecting potential predator ambushes on flowers, bumblebees selectively prolong aerial inspections to boost the accuracy of detection, sacrificing foraging speed to improve survival likelihood [18]. Thus, perceived risks drive context-specific SATs, potentially extending to analogous hazards in route choice.

In this study, we test these spatial perceptual mechanisms in honeybees undertaking an aperture traversal task with two alternatives. Given tight height difference margins, bees’ choices are tracked to explore how they process the visual environment under strict physical constraints.

## Methods

### (a) Experimental setup and procedure

Foragers from a colony of honeybees (*Apis mellifera*) housed in a free-foraging urban apiary on a rooftop at UNSW Canberra were trained to navigate a custom-built wooden tunnel (figure 1; movie S1). The tunnel (0.3 m x 0.3 m x 2.0 m) was positioned approximately 20 m away from the hive on an outdoor rooftop terrace. An overhead shade-sail provided filtered, diffuse natural lighting. The tunnel had a unidirectional design (inlet to outlet) to ensure consistent traffic flow (figure 1*a*). The walls and floor were lined with a grayscale pink noise (1/f) cloud pattern to provide steady OF cues. The ceiling consisted of a 5 mm thick transparent, UV-blocking acrylic sheet to allow illumination and observation of behavior.

**Figure 1.**
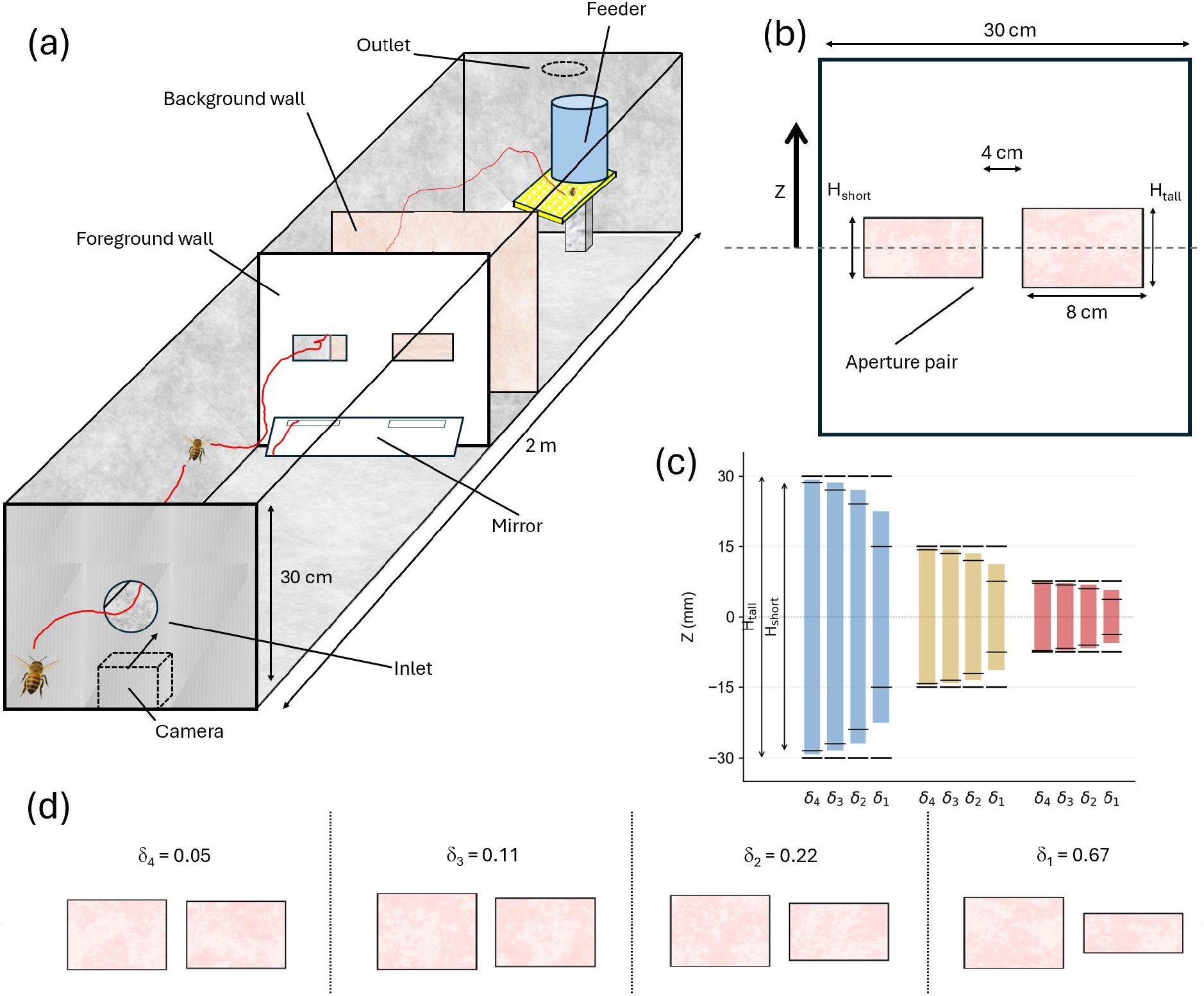
Tunnel configuration and height discrimination stimuli. *(a)* Schematic of the flight tunnel showing the plain foreground wall, textured background wall, and a representative bee trajectory (red line). The camera and its viewing direction is shown below the inlet, along with a 45° mirror at the base of the foreground aperture partition to capture the lateral position of the bee while it chooses between the apertures and immediately before it transits the chosen aperture. *(b)* Front view of the aperture pairs on the foreground wall. The difference in aperture heights is expressed as the relative height difference, 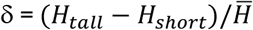 (where *H*_*tall*_ − *H*_*short*_ is the height difference, Δ*H*, and 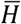 is the mean height. *(c)* Collation of all tested absolute aperture heights adjacent one another in descending order. Thick black horizontal lines indicate the vertical space of the taller aperture (H_tall_), and thin black horizontal lines indicate the vertical space of the shorter aperture (H_short_). The colored bars represent the reference stimulus magnitude (average height, 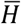) for each pair, grouped by absolute size regime (Blue: Large, Yellow: Medium, Red: Small). *(d)* Frontal view of the four relative height differences (*δ*) as viewed by approaching bees. The apertures are surrounded by an untextured white foreground, and the background noise pattern is visible through the aperture.

An interchangeable transverse partition (0.3 m x 0.3 m), constructed from 80 GSM white paper, was positioned at the midpoint of the tunnel (1 m from the inlet). A shorter background partition (0.3 m x 0.25 m) was placed 20 cm behind the foreground partition. This background had a red-and-white pink noise texture, which would be perceived as high-contrast black-and-white by honeybees because their visible spectrum does not extend to red wavelengths. A camera (GoPro Hero 11, 2K at 60 fps) positioned below the inlet captured the frontal view, using a 45° slanted mirror at the base of the foreground partition to record upward views of bee entries (figure 1*a*).

Bees were trained to fly into the tunnel by placing sucrose-soaked cotton buds at successive waypoints from the hive. Once trained, a gravity feeder containing 10% (v/v) sucrose solution was placed at the far end of the tunnel, under the outlet (figure 1*a*). Bees were then trained to traverse the setup through a single large aperture (12 x 5 cm) in the foreground wall, encouraging rapid, hesitation-free flight. Following this training, bees were tested by presenting them with dual-aperture configurations. Interspersed between tests, continued training bouts with the single large aperture maintained consistent behavior throughout the 3-4 week experimental period.

We estimate that >100 unique foragers routinely commuted through the tunnel based on the high traffic flux (5-10 bees min^-1^) relative to the typical foraging round-trip duration (estimated as <10 min during initial training). During testing, the aperture pair was randomised by a computer program every 5 minutes (see table S1). Camera recording commenced at each reconfiguration. This randomization scheme and the large sample sizes prevented pseudoreplication from repeat visits. Testing occurred on consecutive days during summer (January-February) between 10:00 and 18:00. Each 5-minute interval spanned approximately 10-30 flights, with random swapping continuing until n ≥ 100 flights were captured per condition.

### (b) Stimulus design and data collection

Aperture pairs (figure 1*b*) were carefully cut from 13 separate interchangeable foreground walls. To achieve symmetry, the apertures were positioned with their centers aligned to the vertical center of the tunnel (Z = 0, figure 1*b*) and spaced midway between the horizontal center and the wall. All apertures were 80 mm wide, ensuring an aspect ratio (width/height) well above 1, and featured a plain foreground that confined the approach view to the aperture boundaries and the background wall.

To control for lateral bias, each aperture pair was tested with the taller aperture on both the left (A-L) and right (M-X) by rotating the wall 180° about the longitudinal tunnel axis (all test configurations listed in table S1). Both presentations were subsequently pooled. By varying gap heights while holding other visual cues constant, this design maintained symmetric lateral OF, ensuring selection was based solely on height cues. Although phototaxis could exert a minor influence [12,19], the perceptual resolution of geometrical differences (via OF) likely far exceeds that for brightness stimuli in this configuration. An identical aperture pair (30 mm height) was tested in both default and rotated orientations to confirm the absence of wall orientation effects.

To test the Weber relationship, we applied four proportional height decrements 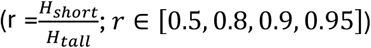 (figure 1*c,d*) across three baseline scales (*H*_*tall*_ = 15, 30, 60 mm). To avoid the normalization bias inherent in using either individual height as the denominator, we used the mean height (*I* = (*H*_*tall*_ + *H*_*short*_)/2) as the reference stimulus magnitude. The relative height difference (RHD) was then derived as:

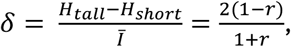

This gives a standardized range of RHD (0 ≤ *δ* ≤ 0.67) across all absolute scales. We manually reviewed sample flights in videos offline, applying a selection process where only direct flights (not those identified as learning flights) were evaluated. Valid choices were tallied until at least 200 flights were recorded per pooled height pair (n = 800-900 per scale, excluding controls).

### (c) Psychometric modelling and statistical analyses

All analyses were conducted in Python (v3.10) using Statsmodels (v0.14) for model fitting, SciPy (v1.15) for statistical tests, and Matplotlib (v3.10) for visualisation. Choice data (c successes out of n trials), were modelled using a four-parameter base psychometric function:

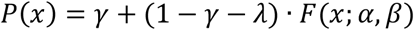

In this general framework, *γ* is the guess rate (defining the lower asymptote), λ is the lapse rate (defining the upper asymptote at 1-λ), and *F*(*x*; *α, β*) is the underlying sigmoid function determined by threshold *α* and slope β, as described below.

We constrained the general function based on our experimental design: we fixed the guess rate to chance level for a 2AFC task (*γ* = 0.5) and assumed no lapses (*λ* = 0), justified by the consistent training and strict manual sample screening process.

The underlying function was modelled using a Weibull distribution, resulting in the following two-parameter model:

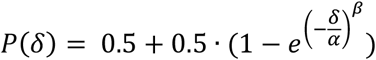

Here, *δ* represents the RHD, *α* is the threshold (stimulus level at which performance reaches ∼81.6% of the dynamic range), and β determines the slope. Model selection via Akiake Information Criterion (AIC) supported this reduced model over variants with free parameters. Fits were performed via maximum likelihood estimation (MLE). We obtained 95% CIs for parameters and curves by performing 5000 bootstrap resamples.

In 2AFC, the Just-Noticeable-Difference (JND) is usually defined as the stimulus difference required to reach P = 75% [20]. However, high behavioral variability and data sparsity in insect studies make it necessary to have lower levels. We selected two thresholds, at 60% and 70%, to robustly compare sensitivity between scales. This allows analysis of asymmetric curves and reduces extrapolation bias while maintaining ecological relevance [21].

Pairwise differences in threshold values and slopes at the 60% and 70% performance levels were evaluated between scales using 5000 bootstrap resamples per scale. Two-tailed empirical p-values were derived from the distributions of differences between the bootstrap samples. Asterisks in figures 2b and 2c indicate significant differences (p*<0.05, p**<0.01). We used Likelihood Ratio Test (LRT) to compare the significance variations between psychometric curves.

**Figure 2.**
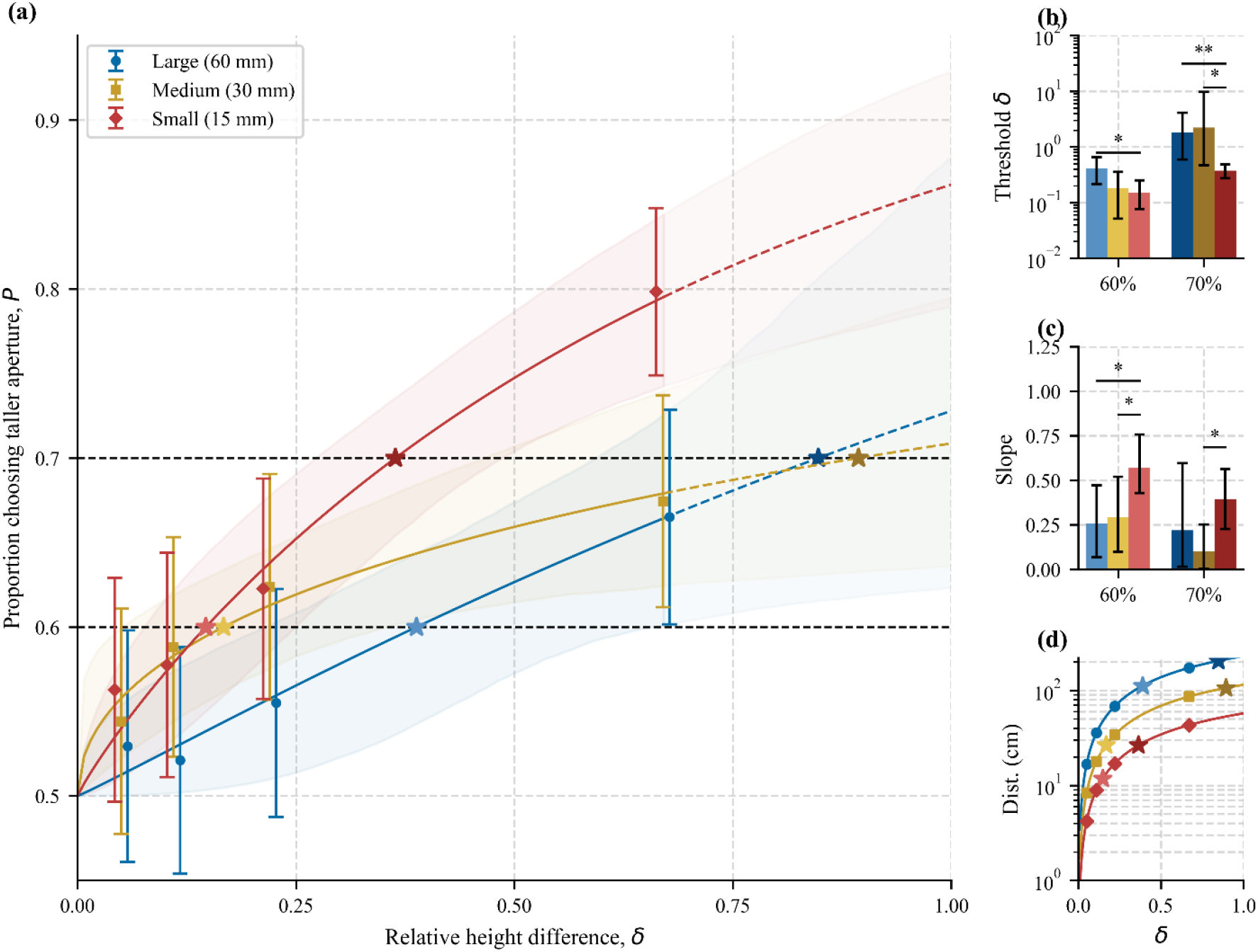
Psychophysics of aperture height discrimination by bees for the three absolute scales. *(a)* Main psychometric functions showing the proportion (P) of bees choosing the taller aperture as a function of relative height difference (*δ*). Data points (mean with 95% binomial CIs) and fitted Weibull curves (solid lines within the tested range, dashed lines in the extrapolated part) with 95% CIs (shaded regions) are shown. Stars indicate thresholds at 60% (light) and 70% (dark) performance levels. *(b)* Threshold relative height differences (*δ*) at 60% and 70% performance thresholds across scales (log scale; means with 95% bootstrap CIs), with asterisks indicating significant pairwise difference (p*<0.05, p**<0.01). *(c)* Slopes at 60% and 70% performance thresholds across scales (means with 95% bootstrap CIs), with pairwise bars and asterisks indicating significant differences. *(d)* Furthest resolvable inspection distance (cm) as a function of *δ*, based on a 1° minimum interommatidial angle. Markers are shown for all tested *δ* and the 60% (light) and 70% (dark) performance thresholds. *Note: Bootstrap CIs are asymmetric (2.5-97.5% percentiles); binomial CIs are Wilson score intervals.

Theoretical maximum inspection distances for resolvability were calculated as:

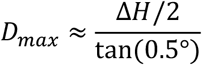

based on a half-interommatidial angle of 0.5° (corresponding to a full interommatidial angle of ∼1° in the frontal eye region [22]). This estimates the farthest distance at which the absolute height difference is resolvable, given angular resolution limits. *D*_60_ values (inspection distance required at the 60% performance threshold) were computed analogously using the absolute height difference corresponding to that threshold (ΔH_60_) and are marked in figure 2*d*.

## Results

Honeybees strongly preferred the taller of the two apertures across all conditions, choosing it in 61% of 2584 flights (binomial test against 50% chance, p < 0.001, Table S2). This preference became more pronounced as the relative height difference between the two apertures increased (figure 2*a*, Tables S2-3).

While discrimination generally followed Weber’s law (performance driven by relative rather than absolute height differences as shown in figure 2*a*), absolute height influenced sensitivity (global LRT p = 0.0013, Table S5). Performance was best at the smallest scale where absolute differences were smaller. At this scale, bees required only modest relative differences (*δ* = 0.15-0.2) to reach 60-70% accuracy (figure 2*b*, table S4). At the large scale, they required a relative difference roughly twice as large to achieve equivalent performance (figure 2*b*). At the medium scale, performance rose sharply at low *δ* but saturated early, reaching a lower maximum accuracy than the other two scales (rapidly levelling yellow curve in figure 2*a*).

Bees were successful at traversing the shortest apertures tested (7.5 mm in the I/U pairs; Table S1, Movie S2), although rare here due to strong preference for the taller gap. Bees generally had flatter body attitudes (more horizontal) (Movie S3), and increased yaw deviations (Movie S4) for these most challenging of gaps. For identical 30 mm tall gaps, bees chose with no significant lateral bias (n = 214; proportion left = 0.53; p = 0.452; figure S1, tables S2, S6). This shows that the observed preferences were driven by visual discrimination of height differences, and not influenced by side bias. Full statistical details of results are provided in the electronic supplementary material, Tables S2-S6.

## Discussion

The behavioural observations suggest that honeybees discriminate between aperture heights primarily based on the relative differences between the apertures. This aligns with Weber’s law, which predicts that discrimination performance depends on the ratio of the stimuli rather than their absolute magnitude [7]. Plotting choice performance against the RHD (figure 2*a*) shows positive monotonic psychometric functions for all scales. Choice probabilities rise from the PSE (P = 0.5) to strong performance at large stimulus magnitudes (figure 2*a*). This broadly consistent rising profile implies that relative difference is the primary driver of discrimination, enabling bees to perform effectively across varying stimulus magnitudes.

However, the choice curves differed significantly across absolute aperture sizes, producing statistically distinct sigmoids (confirmed by LRT as shown in Table S5). This indicates that discrimination involves more than pure relative processing, as the slopes were not ordered by aperture size. Instead, the choice curve for the medium-sized apertures was flatter than both the small and large scales (see reduced slope and elevated thresholds across *δ* in figure 2*a,b,c*). Although effects of absolute size might sometimes be expected (for example, poorer discrimination at small scales due to limited visual resolution [22], or at large scales due to sensory saturation [23]), the results here do not fit such a simple trend. Rather, the particularly shallow curve at medium scales (figure 2*c*) suggests that bees do not simply maximise precision at all times, but instead actively regulate their perceptual effort based on the specific demands and risks of the environment.

The aperture-scale dependent variations shown in the data suggest that choosing between two apertures involves a dynamic risk-cost trade-off, in which collision danger is weighed against the metabolic costs of inspection. This parallels speed-accuracy trade-offs (SATs) in bee decisions, where slower actions enhance accuracy but raise energetic costs [17,24]. While closer, slower approaches enhance geometric acuity [25], they require active deceleration and costly low-speed flight [26]. At small gap sizes, where collision is likely, bees appear to invest in this costly inspection to discern safer passage. This investment appears metabolically affordable because bees must already decelerate to traverse the narrow gap [14]. In contrast, the flattened medium-scale curve (figure 2*a*) could be a transitional zone of relaxed selective pressure. Here, and for larger scales, bees adopt faster, less vigilant flight patterns. This suggests a strategic allocation of attention, where bees accept lower precision to conserve energy.

Similar deviations occur in sugar concentration and numerosity discrimination [11,27], indicating a departure from Weber’s law when excessive precision is not ecologically mandated.

Once committed to one side, the bee must ensure that its body fits the available vertical clearance. We observed bees actively manipulating their posture, flattening their pitch from the typical ∼40° hover to a more horizontal attitude (Movie S3). This can reduce their vertical profile down to the thorax thickness (3.6 mm) [28,29]. For many of the small-scale apertures (<15 mm) this is physically compulsory for traversal, as the standard ∼10 mm vertical profile (at 40°) is physically taller (e.g. for cases I, U) or leaves insufficient free room for variation (cases J-L, V-X; Table S1). Such posturing for safe passage suggests a body-referenced awareness of clearance in the vertical domain and is analogous to the yaw posturing bumblebees use for width-constrained gaps [15]. However, flattening the body limits pitch for rearward force vectoring (directing the mean aerodynamic force backwards). The bees’ increased yaw and sideslip maneuvers observed in the smallest scale (see movie S4) were likely to compensate for this loss of control authority. Bees adopt a sideways orientation to modulate the force vector along the flight path via body roll [30], separating acceleration from attitude. The result is a fixed body pitch that allows trajectory adjustments for maintaining clearance.

We postulate that a risk assessment occurs the instant the visual system can physically resolve the stimulus. The afforded action—to inspect or traverse directly—arbitrages collision safety against the metabolic costs of maneuvering. This mechanism likely complements the Weber principle, facilitating spontaneous, scalable responses calibrated for rapid threading through clutter.

## Electronic supplementary material

**Figure S1**. Analysis of lateral bias across all experimental conditions.

**Table S1**. Geometric specifications of stimuli across all tests.

**Table S2**. Aperture choice results and statistics across all aperture scales and RHDs.

**Table S3**. Psychometric function (Weibull) parameters fitted to choice data for each scale.

**Table S4**. Pairwise statistical comparisons (A vs. B) of performance thresholds and slopes across spatial scales.

**Table S5**. LRT comparisons of psychometric functions across scales.

**Table S6**. Assessment of lateral bias across all experimental conditions.

**Movie S1**. Example approach and traversal through a standard 30 mm aperture pair.

**Movie S2**. Example traversal of a 7.5 mm aperture.

**Movie S3**. Example traversal with pitch adjustment through a narrow aperture.

**Movie S4**. Example traversal with lateral movement through a narrow aperture.

## Acknowledgements

We thank Richard Allum for assistance with laboratory spaces, and Collin for maintaining the apiary.

## References

1. Srinivasan M V. 2011 Visual control of navigation in insects and its relevance for robotics. Curr Opin Neurobiol 21, 535–543.

2. Srinivasan M V. 2021 Vision, perception, navigation and ‘cognition’in honeybees and applications to aerial robotics. Biochem Biophys Res Commun 564, 4–17.

3. Riley JR, Greggers U, Smith AD, Reynolds DR, Menzel R. 2005 The flight paths of honeybees recruited by the waggle dance. Nature 435, 205–207.

4. Burnett NP, Badger MA, Combes SA. 2022 Wind and route choice affect performance of bees flying above versus within a cluttered obstacle field. PLoS One 17, e0265911.

5. Buatois A, Mailly J, Dubois T, Lihoreau M. 2024 A comparative analysis of foraging route development by bumblebees and honey bees. Behav Ecol Sociobiol 78, 8.

6. Worsley MZ, Schroeder J, Dixit T. 2025 How animals discriminate between stimulus magnitudes: a meta-analysis. Behavioral Ecology 36, araf025.

7. Fechner GT. 1966 Elements of Psychophysics. New York, NY: Holt, Rinehart and Winston.

8. Avarguès-Weber A, d’Amaro D, Metzler M, Dyer AG. 2014 Conceptualization of relative size by honeybees. Front Behav Neurosci 8, 80.

9. Hempel de Ibarra N, Vorobyev M, Menzel R. 2014 Mechanisms, functions and ecology of colour vision in the honeybee. Journal of Comparative Physiology A 200, 411–433.

10. Bortot M, Stancher G, Vallortigara G. 2020 Transfer from number to size reveals abstract coding of magnitude in honeybees. iScience 23.

11. Nachev V, Stich KP, Winter Y. 2013 Weber’s law, the magnitude effect and discrimination of sugar concentrations in nectar-feeding animals. PLoS One 8, e74144.

12. Ong M, Bulmer M, Groening J, Srinivasan M V. 2017 Obstacle traversal and route choice in flying honeybees: Evidence for individual handedness. PLoS One 12, e0184343.

13. Ravi S, Bertrand O, Siesenop T, Manz L-S, Doussot C, Fisher A, Egelhaaf M. 2019 Gap perception in bumblebees. Journal of Experimental Biology 222, jeb184135.

14. Jakobi T, Garratt M, Srinivasan M, Ravi S. 2025 How honeybees perceive and traverse apertures. Journal of Experimental Biology, jeb-250145.

15. Ravi S, Siesenop T, Bertrand O, Li L, Doussot C, Warren WH, Combes SA, Egelhaaf M. 2020 Bumblebees perceive the spatial layout of their environment in relation to their body size and form to minimize inflight collisions. Proceedings of the National Academy of Sciences 117, 31494–31499.

16. Chittka L, Skorupski P, Raine NE. 2009 Speed–accuracy tradeoffs in animal decision making. Trends Ecol Evol 24, 400–407.

17. Chittka L, Dyer AG, Bock F, Dornhaus A. 2003 Bees trade off foraging speed for accuracy. Nature 424, 388.

18. Ings TC, Chittka L. 2008 Speed-accuracy tradeoffs and false alarms in bee responses to cryptic predators. Current Biology 18, 1520–1524.

19. Baird E, Dacke M. 2016 Finding the gap: a brightness-based strategy for guidance in cluttered environments. Proceedings of the Royal Society B: Biological Sciences 283, 20152988.

20. Ulrich R, Miller J. 2004 Threshold estimation in two-alternative forced-choice (2AFC) tasks: The Spearman-Kärber method. Percept Psychophys 66, 517–533.

21. Giurfa M, Vorobyev M, Kevan P, Menzel R. 1996 Detection of coloured stimuli by honeybees: minimum visual angles and receptor specific contrasts. Journal of Comparative Physiology A 178, 699–709.

22. Rigosi E, Wiederman SD, O’Carroll DC. 2017 Visual acuity of the honey bee retina and the limits for feature detection. Sci Rep 7, 45972.

23. Spaethe J, Tautz J, Chittka L. 2001 Visual constraints in foraging bumblebees: flower size and color affect search time and flight behavior. Proceedings of the National Academy of Sciences 98, 3898–3903.

24. MaBouDi H, Marshall JAR, Dearden N, Barron AB. 2023 How honey bees make fast and accurate decisions. Elife 12, e86176.

25. Land MF. 1997 Visual acuity in insects. Annu Rev Entomol 42, 147–177.

26. Nachtigall W, Hanauer-Thieser U, Mörz M. 1995 Flight of the honey bee VII: metabolic power versus flight speed relation. Journal of Comparative Physiology B 165, 484–489.

27. Bortot M, Agrillo C, Avarguès-Weber A, Bisazza A, Miletto Petrazzini ME, Giurfa M. 2019 Honeybees use absolute rather than relative numerosity in number discrimination. Biol Lett 15, 20190138.

28. Taylor GJ, Luu T, Ball D, Srinivasan M V. 2013 Vision and air flow combine to streamline flying honeybees. Sci Rep 3, 2614.

29. Winston ML. 1991 The biology of the honey bee. harvard university press.

30. Taylor GK. 2001 Mechanics and aerodynamics of insect flight control. Biological Reviews 76, 449–471.

