## Supplementary data for "Threading the needle: Spatial constraints sharpen visual sensitivity in honeybees"

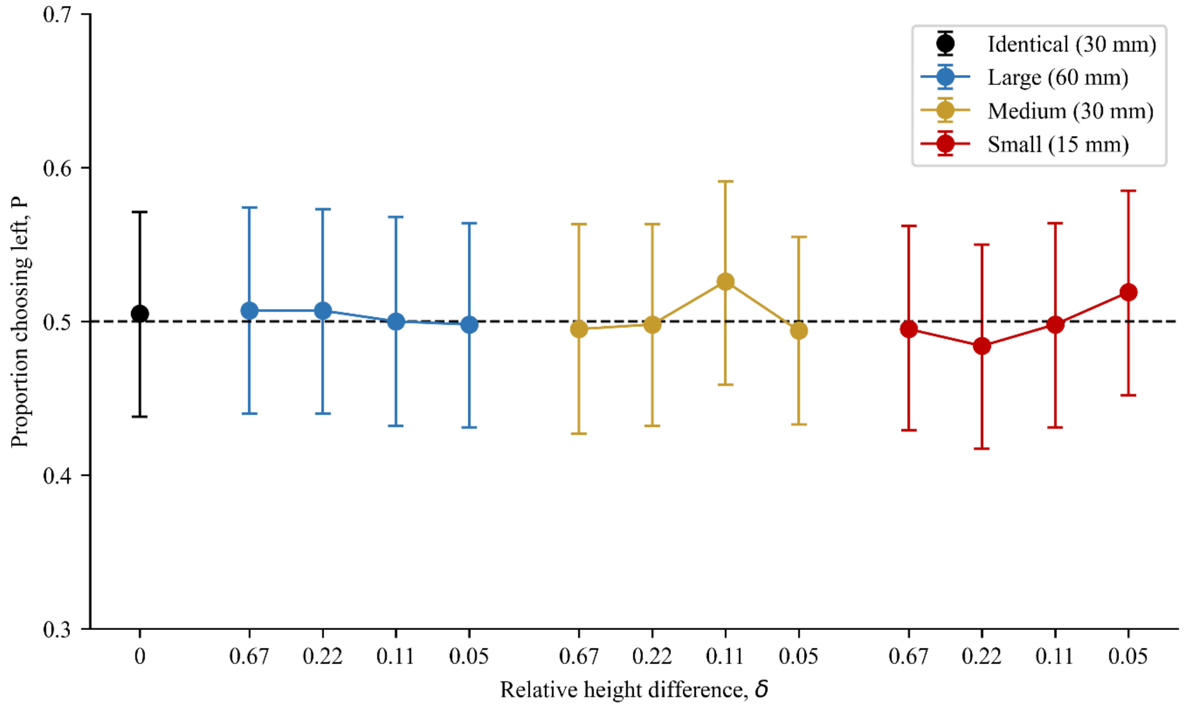

**Figure S1.** Analysis of lateral bias across all experimental conditions.

The proportion (P) of flights choosing the left aperture (after pooling rotated/unrotated sets) is plotted against relative height difference (δ) for each spatial scales. The black marker (δ = 0; identical 30 mm apertures). Error bars are 95% Wilson CIs. The dashed horizontal line represents the chance level (0.5). In all conditions, the CIs overlap the chance line, confirming that the bees displayed no significant lateral bias regardless of aperture scale or difficulty.

**Table S1.** Geometric specifications of stimuli across all tests. Each row details the aperture pair configuration, including the absolute heights of the tall ($H_{tall}$) and short ($H_{short}$) apertures and the side of the taller option. Derived parameters include the ratio r ($H_{short}/H_{tall}$), the absolute height difference $\Delta H$ ($H_{tall} - H_{short}$), the mean height $\bar{H}$ ($(H_{tall} + H_{short})/2$), and the relative height difference δ ($\Delta H/\bar{H}$). Rows Y and Z represent symmetrical control trials (30|30 mm), where Z is a 180° rotation of Y about the longitudinal axis of the tunnel to assess wall orientation bias. Data from unrotated (tall-left) and rotated (tall-right) presentations were pooled for analysis to control for lateral bias.

| Aperture pair | Heights ($H_{tall}$ \| $H_{short}$) | Tall Side | r | ΔH (mm) | $\bar{H}$ (mm) | $\delta$ |
|---|---|---|---|---|---|---|
| A | 60\|30 | | 0.5 | 30 | 45 | 0.67 |
| B | 60\|48 | | 0.8 | 12 | 54 | 0.22 |
| C | 60\|54 | | 0.9 | 6 | 57 | 0.11 |
| D | 60\|57 | | 0.95 | 3 | 58.5 | 0.05 |
| E | 30\|15 | | 0.5 | 15 | 22.5 | 0.67 |
| F | 30\|24 | Left | 0.8 | 6 | 27 | 0.22 |
| G | 30\|27 | (unrotated) | 0.9 | 3 | 28.5 | 0.11 |
| H | 30\|28.5 | | 0.95 | 1.5 | 29.3 | 0.05 |
| I | 15\|7.5 | | 0.5 | 7.5 | 11.3 | 0.67 |
| J | 15\|12 | | 0.8 | 3 | 13.5 | 0.22 |
| K | 15\|13.5 | | 0.9 | 1.5 | 14.3 | 0.11 |
| L | 15\|14.25 | | 0.95 | 0.75 | 14.6 | 0.05 |
| M | 60\|30 | | 0.5 | 30 | 45 | 0.67 |
| N | 60\|48 | | 0.8 | 12 | 54 | 0.22 |
| O | 60\|54 | | 0.9 | 6 | 57 | 0.11 |
| P | 60\|57 | | 0.95 | 3 | 58.5 | 0.05 |
| Q | 30\|15 | | 0.5 | 15 | 22.5 | 0.67 |
| R | 30\|24 | Right | 0.8 | 6 | 27 | 0.22 |
| S | 30\|27 | (rotated) | 0.9 | 3 | 28.5 | 0.11 |
| T | 30\|28.5 | | 0.95 | 1.5 | 29.3 | 0.05 |
| U | 15\|7.5 | | 0.5 | 7.5 | 11.3 | 0.67 |
| V | 15\|12 | | 0.8 | 3 | 13.5 | 0.22 |
| W | 15\|13.5 | | 0.9 | 1.5 | 14.3 | 0.11 |
| X | 15\|14.25 | | 0.95 | 0.75 | 14.6 | 0.05 |

<table>
<tr><td></td><td></td><td></td><td></td><td></td><td></td><td></td></tr>
<tr><td>Y</td><td>30|30</td><td>(unrotated)</td><td>1</td><td>0</td><td>30</td><td>0</td></tr>
<tr><td>Z</td><td>30|30</td><td>(rotated)</td><td>1</td><td>0</td><td>30</td><td>0</td></tr>
</table>

**Table S2.** Aperture choice results and statistics across all aperture scales and relative height differences. Columns show absolute height ratio (r = $H_{short}/H_{tall}$), relative height difference (δ), total sample size (n), and the number of choices for the taller aperture (b). P represents the proportion of correct choices (b/n) with 95% Wilson CIs. P-values (binomial test) represent significance against chance (0.5). Note: The shaded row represents the symmetrical control condition (δ = 0, tested at the medium scale); for this condition, b represents one discreetly marked side tracked across flipped and unflipped tests). Asterisks denote significance: * p<0.05, ** p<0.01, *** p<0.001.

| Scale | r | $\delta$ | n | b | Proportion (P) | 95% CI | p-value |
|---|---|---|---|---|---|---|---|
| Large | 0.50 | 0.67 | 212 | 141 | 0.67 | [0.60, 0.73] | <0.001*** |
| | 0.80 | 0.22 | 209 | 116 | 0.56 | [0.49, 0.62] | 0.128 |
| | 0.90 | 0.11 | 213 | 111 | 0.52 | [0.45, 0.59] | 0.584 |
| | 0.95 | 0.05 | 204 | 108 | 0.53 | [0.46, 0.60] | 0.441 |
| Medium | 0.50 | 0.67 | 215 | 145 | 0.67 | [0.61, 0.73] | <0.001*** |
| | 0.80 | 0.22 | 202 | 126 | 0.62 | [0.56, 0.69] | 0.001** |
| | 0.90 | 0.11 | 221 | 130 | 0.59 | [0.52, 0.65] | 0.010* |
| | 0.95 | 0.05 | 215 | 117 | 0.54 | [0.48, 0.61] | 0.220 |
| | 1 | 0 | 214 | 113 | 0.53 | [0.46, 0.60] | 0.452 |
| Small | 0.50 | 0.67 | 253 | 202 | 0.80 | [0.74, 0.84] | <0.001*** |
| | 0.80 | 0.22 | 212 | 132 | 0.62 | [0.56, 0.69] | <0.001*** |
| | 0.90 | 0.11 | 213 | 123 | 0.58 | [0.51, 0.64] | 0.028* |
| | 0.95 | 0.05 | 215 | 121 | 0.57 | [0.50, 0.63] | 0.076 |

**Table S3.** Psychometric function (Weibull) parameters fitted to choice data for each scale. The model assumes a fixed guess rate ($\gamma = 0.5$) and lapse rate ($\lambda = 0$). $\alpha$ represents the threshold parameter, and $\beta$ represents the shape (slope) parameter. The p-value indicates the empirical probability (from bootstrap resamples) that the slope exceeds zero, showing a significant positive effect of the relative height difference ($\delta$) on choice accuracy.

| Scale | Threshold ($\alpha$) | 95% CI ($\alpha$) | Shape ($\beta$) | 95% CI ($\beta$) | p-value ($\beta>0$) |
|---|---|---|---|---|---|
| Large | 1.6 | [1.02,1.71] | 1.06 | [0.38, 2.5] | <0.001 |
| Medium | 3.48 | [2.30,>100] | 0.49 | [0.19, 0.96] | <0.001 |
| Small | 0.76 | [0.54,1.06] | 0.91 | [0.58, 1.37] | <0.001 |

**Table S4.** Pairwise statistical comparisons (A vs. B) of performance thresholds and slopes across spatial scales. The table evaluates differences in relative height difference ($\delta$) and local psychometric slopes at two performance levels (60% and 70% accuracy). Mean A and Mean B correspond to the first and second groups listed in the Comparison column, respectively. P-values (two-tailed) were derived from 5000 bootstrap resamples. Significance is shown by asterisks: (* $p<0.05$, ** $p<0.01$, *** $p<0.001$).

| Comparison | Mean A (95% CI) | Mean B (95% CI) | p-value |
|---|---|---|---|
| **Threshold at 60% accuracy** | | | |
| Large vs. Medium | 0.41 [0.22, 0.66] | 0.18 [0.05, 0.35] | 0.086 |
| Large vs. Small | 0.41 [0.22, 0.66] | 0.15 [0.08, 0.25] | 0.020* |
| Medium vs. Small | 0.18 [0.05, 0.35] | 0.15 [0.08, 0.25] | 0.809 |
| **Slope at 60% accuracy** | | | |
| Large vs. Medium | 0.26 [0.07, 0.47] | 0.29 [0.1, 0.52] | 0.835 |
| Large vs. Small | 0.26 [0.07, 0.47] | 0.57 [0.43, 0.76] | 0.026* |
| Medium vs. Small | 0.29 [0.1, 0.52] | 0.57 [0.43, 0.76] | 0.047* |
| **Threshold at 70% accuracy** | | | |
| Large vs. Medium | 1.80 [0.59, 4.12] | 2.23 [0.47, 9.74] | 0.998 |
| Large vs. Small | 1.80 [0.59, 4.12] | 0.37 [0.27, 0.49] | <0.001*** |
| Medium vs. Small | 2.23 [0.47, 9.74] | 0.37 [0.27, 0.49] | 0.012* |
| **Slope at 70% accuracy** | | | |
| Large vs. Medium | 0.22 [0.01, 0.6] | 0.10 [0.0, 0.25] | 0.449 |
| Large vs. Small | 0.22 [0.01, 0.6] | 0.39 [0.23, 0.56] | 0.269 |
| Medium vs. Small | 0.10 [0.0, 0.25] | 0.39 [0.23, 0.56] | 0.011* |

**Table S5.** Likelihood Ratio Test (LRT) comparisons of psychometric functions across scales. The table presents the statistical comparison of fitted Weibull models (separate vs. pooled) to assess whether data from different spatial scales can be described by a single underlying function. The Global test compares a model with separate parameters for all three scales against a single pooled model. Pairwise rows compare specific scale pairs. A significant p-value ($p<0.05$) indicates that the psychometric curves are statistically distinct, confirming that absolute aperture size significantly modulates discrimination performance.

| **Comparison** | **LRT statistic ($\chi^2$)** | **df** | **p-value** |
|---|---|---|---|
| Global (all scales) | 17.91 | 4 | 0.0013** |
| Large vs. Medium | 3.54 | 2 | 0.1701 |
| Large vs. Small | 14.35 | 2 | 0.0008*** |
| Medium vs. Small | 8.61 | 2 | 0.0135* |

**Table S6.** Assessment of lateral bias across all experimental conditions. The table shows the raw counts of flights to the physical Left and Right apertures for the symmetrical control (identical) and all asymmetric test conditions (grouped by scale and relative height difference δ). $P_{left}$ is the proportion of choices on the left side (after pooling rotated/unrotated sets). 95% Wilson confidence intervals (CIs) are provided for each proportion. All intervals include the chance level (0.5), suggesting no significant lateral bias in any condition.

| Condition | Left choices | Right choices | Total (n) | $P_{left}$ | 95% CI |
|---|---|---|---|---|---|
| Identical (δ = 0) | 111 | 103 | 214 | 0.519 | [0.452, 0.585] |
| Large (δ = 0.67) | 107 | 105 | 212 | 0.505 | [0.438, 0.571] |
| Large (δ = 0.22) | 106 | 103 | 209 | 0.507 | [0.440, 0.574] |
| Large (δ = 0.11) | 108 | 105 | 213 | 0.507 | [0.440, 0.573] |
| Large (δ = 0.05) | 102 | 102 | 204 | 0.500 | [0.432, 0.568] |
| Medium (δ = 0.67) | 107 | 108 | 215 | 0.498 | [0.431, 0.564] |
| Medium (δ = 0.22) | 100 | 102 | 202 | 0.495 | [0.427, 0.563] |
| Medium (δ = 0.11) | 110 | 111 | 221 | 0.498 | [0.432, 0.563] |
| Medium (δ = 0.05) | 113 | 102 | 215 | 0.526 | [0.459, 0.591] |
| Small (δ = 0.67) | 125 | 128 | 253 | 0.494 | [0.433, 0.555] |
| Small (δ = 0.22) | 105 | 107 | 212 | 0.495 | [0.429, 0.562] |
| Small (δ = 0.11) | 103 | 110 | 213 | 0.484 | [0.417, 0.550] |
| Small (δ = 0.05) | 107 | 108 | 215 | 0.498 | [0.431, 0.564] |
